# L-Aspartate oxidase provides new insights into fumarate reduction in anaerobic darkness in *Synechocystis* sp. PCC6803

**DOI:** 10.1101/2022.10.19.512830

**Authors:** Kateryna Kukil, Jeffrey A. Hawkes, Cecilia Blikstad, Pia Lindberg

## Abstract

Cyanobacteria are promising microbial hosts for production of various industrially relevant compounds, such as succinate, a central metabolite of the tricarboxylic acid cycle (TCA). Cyanobacteria have been engineered to produce succinate during photoautotrophic growth, and are also able to secrete it during anoxic fermentation conditions. It has been assumed that under anoxic darkness, succinate can be formed by reduction of fumarate catalyzed by the succinate dehydrogenase complex (SDH), however, no characterization of SDH regarding this activity has been performed. In this study, we address this issue by generating strains of the unicellular cyanobacterium *Synechocystis* PCC 6803 (*Synechocystis*) deficient in one or several subunits of SDH, and investigating succinate accumulation in these strains during dark anaerobic fermentation. The results showed higher succinate accumulation in SDH deletion strains than in the wild type, indicating a succinate dehydrogenase activity of SDH rather than fumarate reduction under these conditions. We further explored the possibility of another potential route for succinate formation from fumarate via L-aspartate oxidase (Laspo). The gene encoding Laspo in *Synechocystis* could not be inactivated, indicating an essential function for this enzyme. Using purified *Syn*Laspo, we could demonstrate *in vitro* that in addition to L-aspartate oxidation the enzyme exhibits an L-aspartate-fumarate oxidoreductase activity. We therefore suggest that reduction of fumarate to succinate during anoxic darkness can be a byproduct of the Laspo reaction, which is the first step in biosynthesis of NAD cofactors. This work contributes to the understanding of cyanobacterial TCA cycle for future engineering and sustainable production of dicarboxylic acids.

## Introduction

Cyanobacteria have emerged as promising chassis as microbial cell factories for the production of various industrially relevant chemicals by direct carbon capture and conversion. Among those, C4 dicarboxylic acids such as succinate and fumarate are important platform chemicals in the pharmaceutical and food industries [1, 2]. Engineering strategies for succinate production in cyanobacteria have been based on exploiting the oxidative [3–7] or reductive branches of the tricarboxylic cycle (TCA cycle) [8–12].

In light conditions, the TCA cycle in cyanobacteria functions as a branched chain pathway rather than a cycle, synthetizing carbon skeletons for nitrogen assimilation metabolism from 2-oxoglutarate (2-OG) and providing substrate for synthesis of amino acids from oxaloacetate (Figure 1). Succinate can be formed from 2-OG via the γ-aminobutyric (GABA) shunt [13] or via 2-oxoglutarate decarboxylase and succinic semialdehyde dehydrogenase [14]. These bypasses were identified in *Synechocystis* sp. PCC 6803 (hereafter *Synechocystis*) and are closing the cyanobacterial TCA cycle (Figure 1). However, the metabolic flux of carbon from 2-OG to succinate through these bypasses is proposed to be minimal [15].

**Figure 1.**
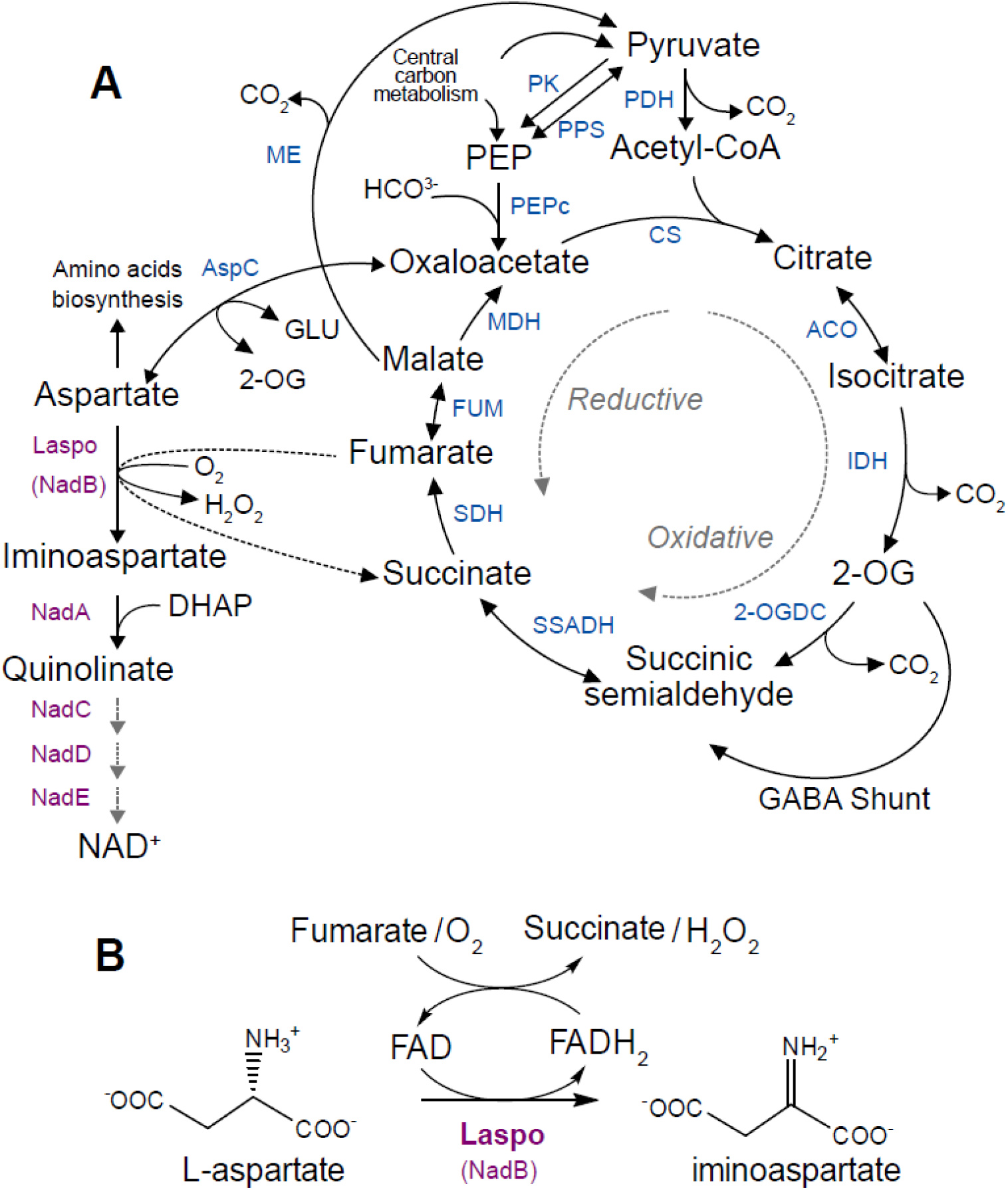
**A** Carbon metabolism involved in the TCA cycle and NAD^+^ cofactor biosynthesis in *Synechocystis* PCC 6803. Abbreviations: PEP, phosphoenolpyruvate; 2-OG, 2-oxoglutarate; 2-OGDC, 2-oxoglutarate decarboxylase; ACO, aconitase; CS, citrate synthase; FUM, fumarase; ACO, aconitase; CS, citrate synthase; FUM, fumarase; GABA, γ-aminobutyrate; IDH, isocitrate dehydrogenase; MDH, malate dehydrogenase; ME, malic enzyme; PDH, pyruvate dehydrogenase complex; PEP, phosphoenolpyruvate; PEPc, phosphoenolpyruvate carboxylase; PK, pyruvate kinase; PPS, phosphoenolpyruvate synthase; SDH, succinate dehydrogenase; SSA, succinic semialdehyde; SSADH, succinic semialdehyde dehydrogenase. **B** Scheme of the reaction catalyzed by Laspo enzyme. Laspo (encoded by *nadB* gene) oxidases L-aspartate to iminoaspartate using either oxygen or fumarate as electron acceptor for FAD reoxidation and forming oxygen peroxide or succinate correspondingly.

In dark conditions and the presence of oxygen, the membrane associated enzyme complex succinate dehydrogenase (SDH) contributes to the reduction of electron transport chain, with concomitant oxidation of succinate to form fumarate [16] and thereby the oxidative branch of TCA cycle is most active. On the other hand, the ability of *Synechocystis* to excrete succinate during dark anoxic fermentation via the reductive branch of the TCA cycle has been explored in several reports [8, 9, 12, 17]. In this pathway, succinate would presumably be formed via reduction of oxaloacetate to form malate and fumarate and finally succinate, by the enzymes malate dehydrogenase, fumarate hydratase and SDH respectively. Based on similarity with other bacteria, such as *E. coli*, succinate formation has been assumed to occur via fumarate reduction by SDH. However, detailed biochemical information on this enzyme complex is absent in cyanobacteria and no fumarate reductase activity has been demonstrated. So far, studies of SDH were carried out by deletion of open reading frames of SDH subunits in *Synechocystis* and subsequent measurements of succinate or fumarate accumulation [3, 18]. While the formation of succinate from fumarate is supported by data from metabolite turnover flux analysis [8], Zhang and co-workers [19] could show that a quadruple knock out mutant in *Synechococcus* PCC 7002 lacking all enzymes known to form succinate (namely SDH, succinyl-CoA synthase, succinic semialdehyde dehydrogenase and 2-oxoglutarate decarboxylase) accumulated similar levels of succinate as the wild type strain.

These reports in the literature raises the question of the existence of another route to synthesize succinate from fumarate. As was proposed previously [19, 20], in cyanobacteria the reduction of fumarate to succinate could occur by activity of the enzyme <u>L</u>-<u>asp</u>artate <u>o</u>xidase (Laspo). In *E. coli*, specifically, Laspo (Uniprot entry P10902) can oxidase aspartate using oxygen or fumarate as electron acceptors forming iminoaspartate and peroxide or succinate, respectively (Figure 1B) [21].

The Laspo reaction is the first step in *de novo* synthesis of NAD cofactors from L-aspartate. In *Synechocystis*, there exist three orthologues of the genes in this pathway: L-aspartate oxidase (EC 1.4.3.16; *nadB; sll0631*), quinolinate synthetase (EC 4.1.99; *nadA; sll0622*), and quinolinate phosphoribosyl-transferase (decarboxylating) (EC 2.4.2.19; *nadC; slr0936*) [22]. Findings in *E. coli* show that the preferable choice of substrate for Laspo is based on the intracellular concentration of fumarate [23]. Fumarate is used as the electron acceptor under anaerobic conditions, whereas under aerobic conditions oxygen is used as electron acceptor. Since the genes encoding the NAD cofactor synthesis pathway is present in *Synechocystis* it is possible that secretion of succinate during anaerobic fermentation can be explained by increased fumarate concentration [12] and Laspo activity.

In this study, we intended to clarify Laspo’s contribution to succinate formation in *Synechocystis*. For this, we created several *Synechocystis* strains with the deletions in SDH open reading frames and *nadB*, and compared their succinate accumulation titers. Furthermore, we purified Laspo from *Synechocystis* (*Syn*Laspo) and measured L-aspartate:fumarate oxidoreductase activity *in vitro*. Our findings provide insights into fumarate reduction in *Synechocystis* during anoxic darkness, which has so far been assumed to be performed by the reversible action of SDH.

## Results

### Succinate accumulation in engineered strains

In this study, we aimed to investigate succinate production during anoxic fermentation, and therefore constructed *Synechocystis* strains where genes encoding SDH or *Syn*Laspo are inactivated. If either enzyme possesses fumarate reductase activity, differences in corresponding knock out strain variants compared to the wild type can be detected. In *Synechocystis*, three ORFs of the SDH complex have been identified, where *sll1625* and *sll0823* encodes two homologous iron-sulfur cluster proteins (subunit B, *sdhBs*) and *slr1233* encodes a flavoprotein (subunit A, *sdhA*) [3, 18]. We created three strains with different SDH subunits knocked out (see Table 1 and Figure 2A): strain SynΔ1625Δ0823 is lacking both B subunits, strain SynΔ1233 is lacking subunit A, and SynΔ1625 strain is deficient in one of the B subunits, deletion of which was previously shown to result in enhanced succinate accumulation during late stationary phase of photoautotrophic growth [3].

**Figure 2.**
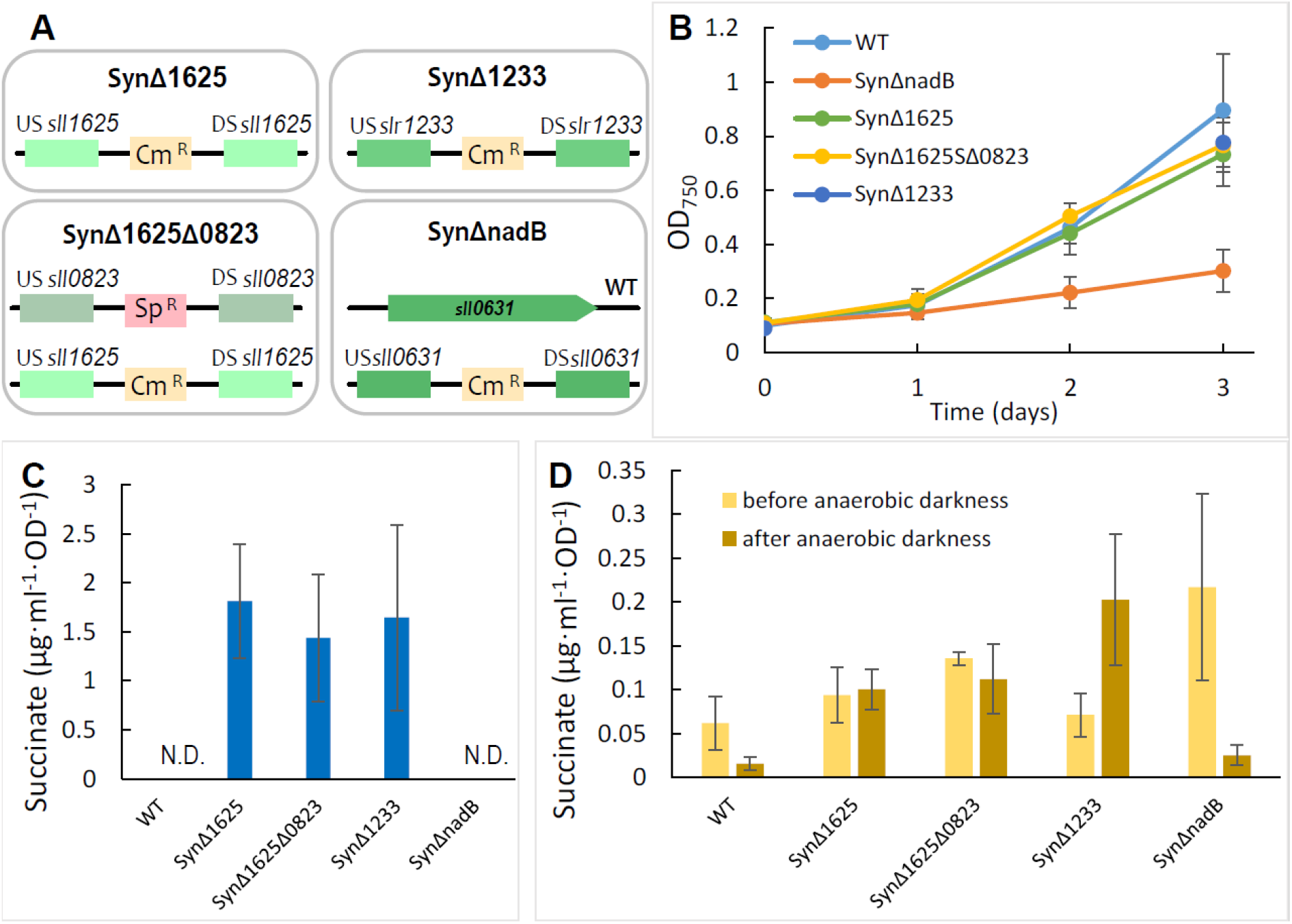
Schematic overview of engineered *Synechocystis* PCC 6803 strains, their growth and succinate levels. **A** Illustration of knockout strains of *Synechocystis* created in this study. **B** Growth curves of knockout strains during three days of cultivation. **C** Succinate accumulated in the growth medium after two days of anoxic fermentation. **D** Intracellular succinate detected before and after anoxic fermentation. OD_750_ refers to optical density at 750 nm. Error bars represent the standard deviation of three biological replicates.

**Table 1.** List of plasmid backbones and *Synechocystis* strains used in this study.

| Construct | Comment | Antibiotics resistance | Reference |
| --- | --- | --- | --- |
| **pEERM3** | Empty backbone based on pJ344 vector backbone for integration into neutral site 1 (*slr1068*) of *Synechocystis* 6803 | Cm | [24] |
| **pEERM3Sp** | Same as pEERM3 but with Sp<sup>R</sup> cassette instead of Cm<sup>R</sup> | Sp | [25] |
| **pET-11a** | Bacterial expression vector based T7 RNA polymerase promotor | Amp | Novagen |
| **p1625** | same as pEERM3 but with homologous recombination regions to *sll1625* | Cm | this study |
| **p0823** | same as pEERM3Sp but with homologous recombination regions to *sll0823* | Sp | this study |
| **p1233** | same as pEERM3 but with homologous recombination regions to *slr1233* | Cm | this study |
| **pnadB** | same as pEERM3 but with homologous recombination regions to *sll0631* | Cm | this study |
| **pET-nadB** | Vector based on pET-11a carrying *SynLaspo* gene *nadB* for overexpression | Amp | this study |
| Strain | Comment | Antibiotics resistance | Reference |
| **SynΔ1625** | *Synechocystis* with disrupted *sll1625* ORF by introducing Cm<sup>R</sup> cassette | **Cm** | this study |
| **SynΔ1625Δ0823** | Same as SynΔ1625, but with disrupted *sll0823* ORF replaced with Sp<sup>R</sup> cassette | **Cm, Sp** | this study |
| **SynΔ1233** | *Synechocystis* with disrupted *slr1233* ORF by introducing Cm<sup>R</sup> cassette | **Cm** | this study |
| **SynΔnadB** | *Synechocystis* with partially disrupted *sll0631* ORF by introducing Cm<sup>R</sup> cassette | **Cm** | this study |

Laspo is a flavoprotein, its structure is homologous to fumarate reductase and SDH, but it lacks their iron-sulfur clusters and membrane attachment subunits [26]. We attempted to inactivate *nadB (sll0631*), encoding Laspo in *Synechocystis*. However, we did not succeed in creating a fully segregated knock out strain, indicating an essential role of this enzyme, as could be expected based on the established role of Laspo in NAD cofactor biosynthesis in other organisms. Nonetheless, the partial knock out strain SynΔnadB was further used in this study.

To investigate the succinate accumulation during anoxic fermentation, obtained strains together with WT as a control strain were grown in shake flasks under photoautotrophic conditions for three days. Even though full segregation was not achieved for the SynΔnadB strain, the culture showed slower growth compared to other knockout strains with same antibiotic cassette resistance (Figure 2B), likely due to effects of the partial gene inactivation. After three days of cultivation, the cultures were concentrated about eight times, transferred to closed vials and flushed with nitrogen gas to induce anaerobicity. Vials were thereafter kept in darkness for two days. The levels of succinate accumulated in the supernatant and intracellularly after two days of anaerobic darkness are presented in Figure 2C and D, respectively.

The SynΔnadB strain behaved as the WT control and no succinate secretion was detected from either strain. All strains lacking SDH subunits secreted higher amounts of succinate than WT or SynΔnadB, however there were no significant differences among these strains (Figure 2C).

Intracellular levels of succinate in the ΔSDH strains after anoxic dark conditions were higher than in the WT and SynΔnadB strains, in agreement with the results on secreted succinate (Figure 2C and D). SynΔnadB exhibited higher intracellular succinate after the photoautotrophic growth period compared to the other strains, but this decreased to WT levels after anoxic darkness incubation (Figure 2D). Although the SynΔnadB strain retains WT copies of *nadB*, it also showed slower growth rate than WT. This growth inhibitory effect might be related to altered fluxes of TCA cycle metabolites due to lower gene dosage of *nadB* in the SynΔnadB strain or an inability to sufficiently provide the required cofactor pool during photoautotrophic growth.

### Properties of purified *Syn*Laspo

Strep-tagged Laspo was expressed in *E. coli* and successfully purified using Strep-Tactin affinity chromatography. The expected size of 62 kDa (including strep-tag) was confirmed by SDS-page and the protein was judged to be over 90% pure (Figure 3). The purified protein fraction was yellow, indicating an oxidized FAD cofactor bound to the protein. Presence of 10% glycerol in the purification buffers was crucial to retain the cofactor [27, 28], as the absence of glycerol resulted in non-colored fraction of inactive enzyme, an indication that the FAD was not bound to the protein.

**Figure 3.**
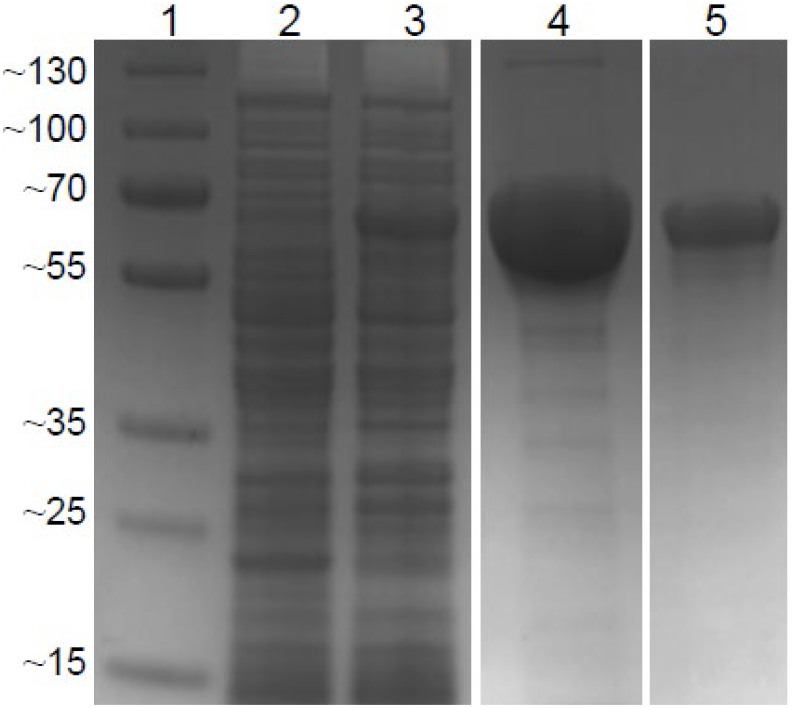
SDS–PAGE analysis showing expression and purification of *Syn*Laspo. Lane 1: molecular weight markers with sizes in kDa. Lane 2: non-induced cells of *E. coli* BL21 (DE3) expressing Laspo (*nadB* gene). Lane 3: induced cells of *E. coli* BL21 (DE3) expressing *Syn*Laspo protein. Lane 4: representative fraction of *Syn*Laspo peak eluted from Strep-Tactin affinity chromatography. Lane 5: diluted fraction of *Syn*Laspo peak. Size of the eluted protein correspond to Laspo’s MW of ~60 kDa and protein were judged to be over 90% pure.

The purified enzyme showed an absorbance spectrum in the visible range typical for FAD– containing flavoproteins (Figure 4A), with two slightly shifted absorbance peaks (375 nm instead of 385 nm and 450nm instead of 445 nm) [29]. An addition of aliquots of L-aspartate and fumarate, the suggested substrates [27], changes the visible spectrum of *Syn*Laspo (Figure 4A): fumarate binding results in hypochromic shift, whereas L-aspartate shifts the maximum absorbance to 463 nm. Anaerobic addition of L-aspartate converts the oxidized FAD into the reduced flavin, which can be observed by disappearance of the absorbance band at 460 nm (Figure 4B). The exposure of the reduced protein to air immediately reoxidizes flavin.

**Figure 4.**
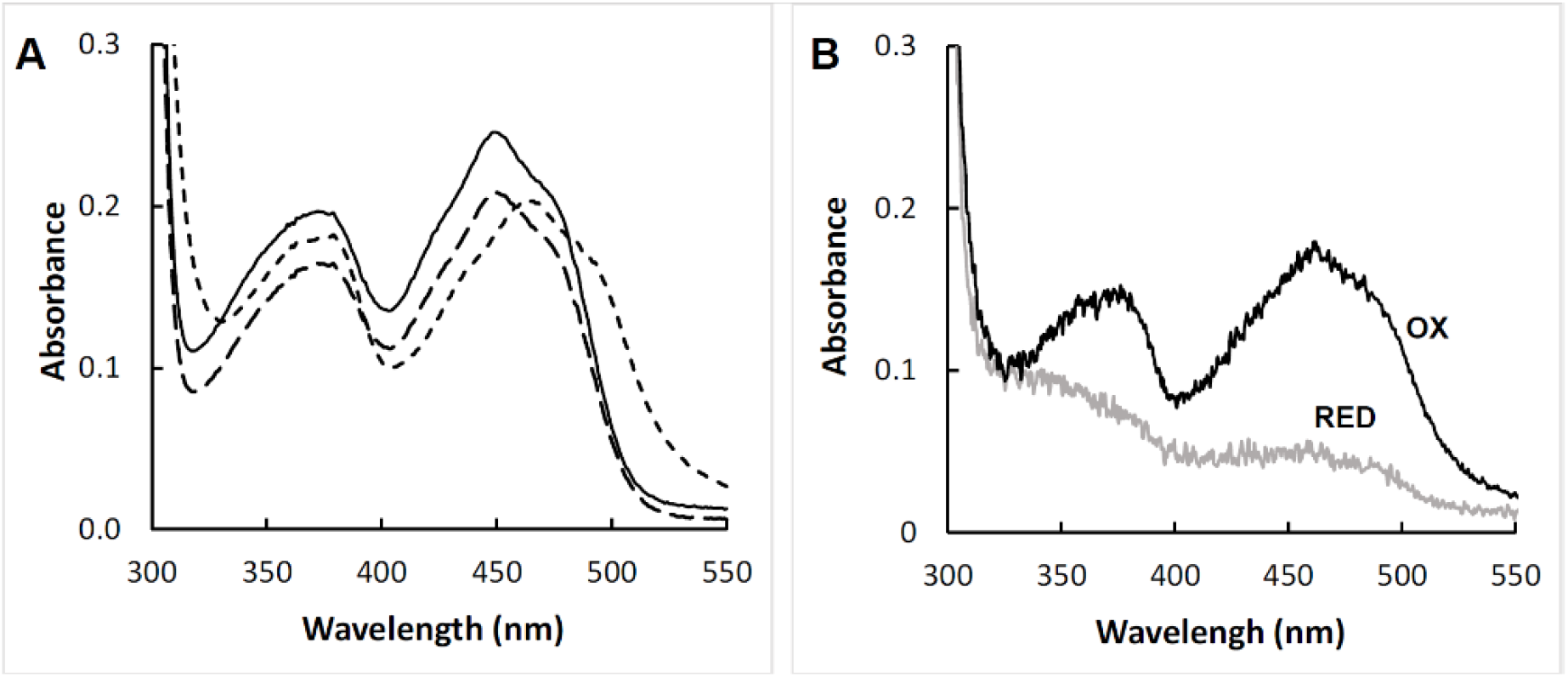
Spectral properties of *Syn*Laspo. **A**: visible absorption of ~50 μM purified *Syn*Laspo in 50 mM potassium phosphate buffer (pH 8) containing 10% glycerol before (---) and after addition of small aliquots of 10 mM L-aspartate (---) and 2 mM fumarate (---). **B**: visible absorption of *Syn*Laspo in different redox states. Absorbance spectrum of oxidized (OX, black line) and reduced (RED, grey line) form of *Syn*Laspo after adding aliquots of 40 mM L-aspartate stock under anaerobic conditions.

The activity of *Syn*Laspo was determined as described in experimental procedures. Briefly, the product of the Laspo reaction, iminoaspartate (see scheme in Figure 2B), is unstable in solution and hydrolyses spontaneously to oxaloacetate and ammonia. We therefore measured oxaloacetate, which can be detected by an established 2,4-dinitrophenylhydrazine (DNPH) coupled assay [30]. As described earlier for *E. coli* Laspo, the catalytic cycle consists of a first half reaction, where FAD is reduced while L-aspartate oxidizes to iminoaspartate and a second half reaction in which FAD is reoxidized using the second substrate (fumarate or O_2_) as electron acceptor (Figure 1B) [31]. However, in our assay tricholoacetic acid is used to quench the reaction by denaturing the protein, making it impossible to conclude whether the product iminoasparte remains bound to the protein or releases from the product-enzyme complex before binding of the second substrate. Laspo oxidoreductase activity was determined by steady-state kinetics by varying L-aspartate concentration. For oxidation of L-aspartate by *Syn*Laspo using oxygen as an electron acceptor, *K*_m_ (the apparent dissociation constant) was measured to 10.2 ± 1 mM and, *k*_cat_ (the turnover number) to 19.2 ± 1 min^-1^. Moreover, substrate inhibition occurred with non-physiological L-aspartate concentration above 30 mM (Figure 5A), which has also been observed for plant Laspo [32].

**Figure 5.**
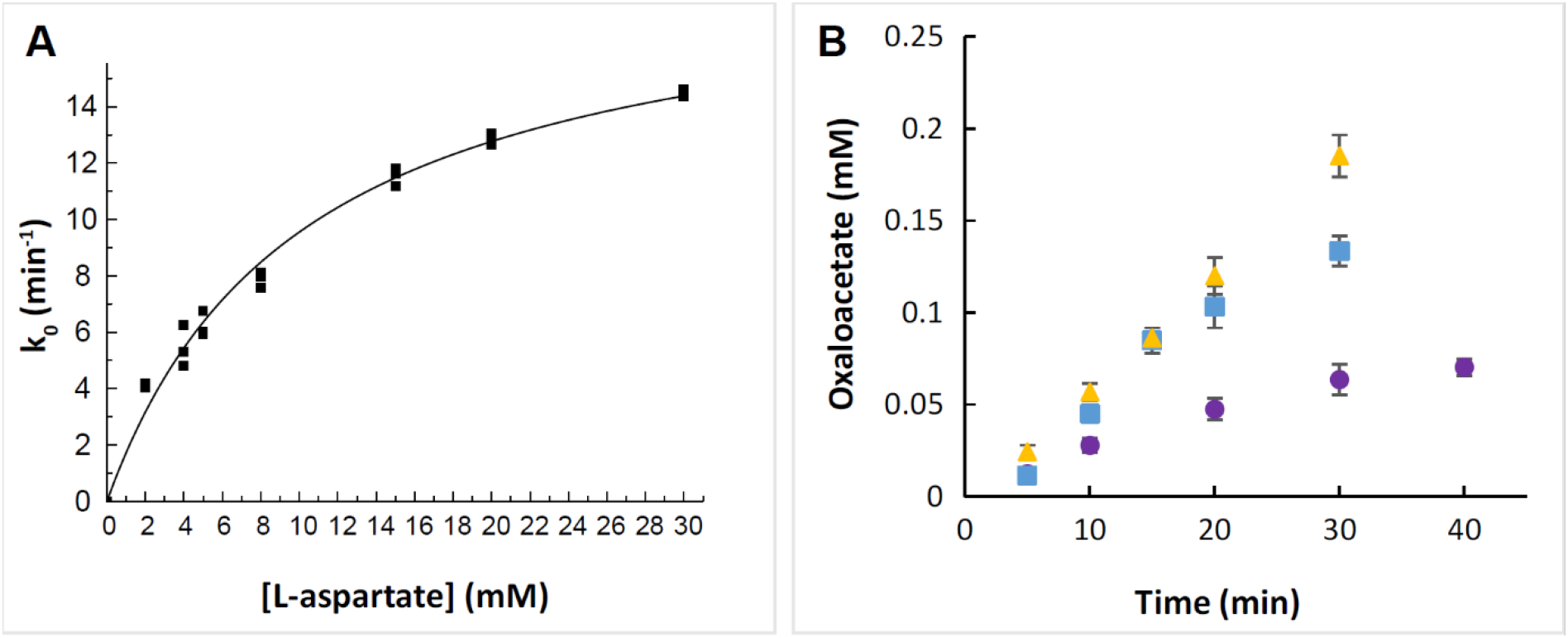
Activity of *Syn*Laspo under aerobic (**A**) and anaerobic conditions (**B**), detecting oxaloacetate formation. **A** Michaelis-Menten fitting of L-aspartate oxidase activity, assayed with varied concentrations of L-aspartate in the reaction mix. Fitting was done using OriginLab software. **B** L-aspartate-fumarate oxidoreductase activity, measured with 20 mM L-aspartate without fumarate 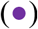, with 0.1 mM fumarate 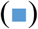 and with 0.5 mM fumarate 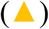 at 30 °C.

The ability of *Syn*Laspo to use fumarate as electron acceptor during oxidation of L-aspartate (Figure 1B) was tested under anaerobic conditions using same experimental assay. As shown in Figure 5B (and Figure 4B) addition of only L-aspartate without the second substrate results in product formation and FAD reduction. This was observed by a decreasing intensity of yellow color of the reaction mix. The presence of fumarate in the reaction mix resulted in increased product formation over time-course of the assay. The appearance of succinate in the reaction mix was confirmed by LC-MS (Figure S1). Taken together, this data demonstrates L-aspartate-fumarate oxidoreductase activity of *Syn*Laspo, in which fumarate can replace oxygen as an electron acceptor.

## Discussion

In this work, we aimed to elucidate the enzymatic route of succinate formation in the model cyanobacterium *Synechocystis* during conditions of anoxygenic fermentation. The reduction of fumarate to succinate through SDH complex was addressed by deletion of SDH subunits. One of the two homologous iron-sulfur proteins, encoded by *sll0823*, has been proposed to be responsible for the fumarate reduction [12], as only the other similar subunit, encoded by *sll1625*, seems to be required for the oxidation of succinate to fumarate [3].

In our experiments, we observed elevated succinate accumulation in SDH deletion strains after anaerobic fermentation, showing no difference between SynΔ1625Δ0823 and SynΔ1625 strains. If the subunit encoded by *sll0823* was responsible for succinate formation under these conditions, a lower succinate concentration would be expected in the double knock-out strain compared to the others. For WT *Synechocystis*, the amounts of succinate obtained in our study after incubation under anaerobic darkness conditions are lower than what was previously reported after anoxic fermentation [8, 9]. This is due to the use of different experimental conditions; using less biomass for anaerobic fermentation, we aimed to avoid the effect cell death, which appears in very dense cultures, resulting in a lower volumetric accumulation of succinate in our experiments.

Previously, it was shown that deletion of SDH subunits alters the glycogen metabolism and assimilation [3]. Glycogen accumulation was increased in SDH knock out strains growing photoautotrophically, possibly due to disrupted regulatory mechanisms of carbon partitioning through the TCA cycle and thus resulting in increased NADH formation, where glycogen would serve as an electron sink [3]. Unlike in photoautotrophic growth, during anaerobic darkness conditions cyanobacteria maintain their viability by breaking down storage carbohydrates. Thus, the elevated succinate concentration in the growth media in our experiments with *Synechocystis* SDH deletion strains under anaerobic darkness may indicate one or several of the following scenarios: (i) increased glycogen accumulation during the first photoautotrophic growth period, due to inactivation of SDH, subsequently leads to a higher succinate secretion due to breakdown of glycogen during the dark anaerobic phase; (ii) succinate formation is not dependent on SDH activity, but instead on other routes such as Laspo activity; (iii) succinate may form during anaerobic fermentation via the oxidative branch of TCA cycle. To the best of our knowledge, an SDH knock out strain of *Synechocystis* has not been investigated in anaerobic conditions via measurements of TCA cycle intermediates. Addressing the SDH properties via *in vivo* succinate or fumarate measurements might be inconclusive due to the plasticity of cellular processes and carbon metabolism, since any changes affecting central carbon metabolism lead to a cascade of alterations involving regulating mechanisms.

As indicated in (ii) above, one candidate other than SDH for succinate formation via fumarate reduction in *Synechocystis* is the Laspo enzyme, which catalyzes the first reaction step in *de novo* biosynthesis of NAD. The final products of NAD biosynthetic pathway, pyridine nucleotides (NAD(P)(H)) participate as cofactors in multiple core redox reactions and have essential cellular functions. NADP^+^, in particular, is a final electron acceptor of photosynthetic electron transport [33]. Thereby, the intracellular concentration of NAD should be tightly regulated according to the cellular needs, which can be achieved by *de novo* synthesis or in recycling (salvaging) pathways. For instance, it was recently shown that the final enzyme in the NAD biosynthesis pathway, NadE, is under global control of carbon/nitrogen levels via PII protein signaling [34] and its activity is suppressed by low nitrogen levels when cell growth is arrested.

In this study we expressed and purified recombinant Laspo enzyme from *Synechocystis* in its active form. Oxygen dependent activity of the enzyme showed a high *K*_M_ for aspartate (10.1±0.6 mM), which is unlike that of other bacterial Laspos from e.g. *E. coli* (1.7 mM) [21, 27], *B. subtilis* (1.0 mM) [35], or the thermophilic archea *Sulfolobus tokodaii* (1.3 mM) [28]. We could also demonstrate the L-aspartate fumarate oxidoreductase activity of the enzyme, showing its ability to function under both aerobic and anaerobic conditions. Therefore, we suggest that succinate may form by Laspo using fumarate as an electron acceptor from aspartate oxidation when the intracellular conditions are switched to anaerobic. This reaction represents an interlink between the TCA cycle intermediates and biosynthesis of essential cofactors. This is in agreement with the study of Cooley and Vermaas 2001 [16], where the total levels of NAD(H) and NADP(H) were much lower in an SDH deficient mutant compared to WT; in absence of SDH activity, fumarate levels would be lower and this may lead to a lack of substrate for the Laspo reaction. The previously reported upregulation of aspartate transferase (*sll0938*) and downregulation of argininosuccinate synthase (*slr0585*), the first step of cyanophycin synthesis from arginine and aspartate, [8] under dark anoxic conditions further support our suggestion.

Although Laspo might use aspartate and molecular oxygen instead of fumarate for catalysis, it is noteworthy that the second enzyme of the NAD biosynthesis, NadA (see Figure 1) may be oxygen sensitive in *Synechocystis*, based on its protein sequence [22, 35]. As described for purified NadA from other organisms [35–37], the iron-sulfur cofactor of this protein is required for the catalysis and is highly oxygen sensitive. Moreover, it has been suggested that the two proteins Laspo (NadB) and NadA form a reversible multienzyme complex for catalysis, which prevents the hydrolysis of unstable iminoaspartate [35], implying that these two enzymes are operating simultaneously. It is interesting thereby, to consider the functionality of the *de novo* NAD biosynthesis in cyanobacteria that are actively producing oxygen during photosynthesis. Although in nature cyanobacteria can experience a decrease of oxygen during dark periods, they are capable of growth under continuous light. Possibly, an oxygen protection mechanism may be involved to ensure the catalysis of these first steps of NAD cofactors biosynthesis. Further studies will be required in order to address this particular issue.

To conclude, in this study we purified Laspo from *Synechocystis* and for the first time demonstrated that *Syn*Laspo is functional under anaerobic conditions. It reduces fumarate to succinate during oxidation of aspartate, the first step in biosynthesis of NAD cofactors. Furthermore, we showed that strains deficient in one or several subunits of SDH accumulate and secrete more succinate than the WT strain during dark anaerobic fermentation, supporting the notion that SDH primarily acts as an SDH and not as a fumarate reductase under these conditions. Biochemical characterization of the cyanobacterial SDH complex is needed to fully address its possible fumarate reductase activity, as deletion of SDH subunits has been shown to lead to multiple primary and secondary effects on cell carbon metabolism. The findings reported here advance our understanding of cyanobacterial TCA cycle metabolism, and will be useful for promoting the development of cyanobacteria as microbial cell factories for sustainable production of high value products such as dicarboxylic acids succinate and fumarate.

## Methods

### Bacterial strains and growth conditions

*Escherichia coli* DH5α (Invitrogen) strain was used subcloning and BL21(DE3) (Thermofisher) for recombinant protein purification. *E. coli* cells were grown in LB medium at 37°C unless other is specified and supplemented with appropriate antibiotics to the final concentrations in the medium: 50 μg ml^-1^ spectinomycin (Sp) or/and 35 μg ml^-1^ chloramphenicol (Cm) or 100 μg·ml^-1^ ampicillin (Sigma, Merk).

*Synechocystis* sp. PCC 6803, a unicellular glucose-tolerant strain was used in this study. Cultures were grown in BG11 medium [38] with respective antibiotics Sp 25 μg ml^-1^ and/or Cm 20 μg ml^-1^ at 30 °C under constant light. The optical density of *Synechocystis* cultures was determined at 750 nm using a Varian Cary 50 BIO spectrophotometer, and an OD_750_ value of 1 corresponds to approximately 10^8^ cells.

### Construction of knockout plasmids and transformation of *Synechocystis*

Deletion of SDH subunits encoded by open reading frames *sll1625, sll0823* and *slr1233* and deletion of *Syn*Laspo encoded by *nadB* gene (*sll0631*) was performed by substitution the corresponding genes with antibiotic resistance cassettes via homologous recombination. For this, 1kb flanking regions upstream and downstream of the coding sequences were amplified from *Synechocystis* genomic DNA and cloned into integrative vectors pEERM3 [24] with Cm resistance creating plasmids p1625, p1233 and pNadB. Similarly, p0823 was constructed based on pEERM3 with Sp resistance (see Table 1.).

For transformation, *Synechocystis* WT cells were transformed as previously described [39]. Colonies that appeared after 10-14 days were checked by PCR for full segregation and restreaked until fully segregated. In order to create the double Δ*sll1625* Δ*sll0823* SDH knockout, the fully segregated *Synechocystis* strain SynΔ1625 was used for transformation with p0823 plasmid.

### Determination of extra- and intracellular succinate concentrations

To assess the levels of succinate during anoxic darkness in the created knockout *Synechocystis* strains, cells were grown in shake flasks with appropriate antibiotics under a constant light intensity of 45 μmol photons m^-2^ s^-1^ from starting OD_750_ ~0.1. After three days, 25 ml of culture were centrifuged and washed twice with fresh BG11 medium, concentrated into 3 ml, and placed into 6 mL headspace vial (Sigma Aldrich). 200 μL of culture sample was taken for metabolites extraction before vials were sealed. The vials were made anaerobic by flushing with nitrogen gas, wrapped with aluminum foil and incubated at 30°C under constant shaking. After two days, 2 ml of culture was used for metabolite extraction, whereas rest was used to measure OD_750_ and centrifuged at 13000 rpm for 10 min at room temperature. The supernatant was stored at −20°C until use.

Extraction of intracellular succinate was performed as described in Prasannan et al [40] with some modifications. Samples of 2 ml of culture collected after anaerobic darkness incubation were centrifuged at 5000 rpm and washed twice with 1ml of 20 mM NaCl. Then 1 ml of ice cold 100% methanol was added to quench the metabolism. The mixture was incubated at −80°C for at least one hour; 1.5 ml of chloroform was added to the extraction mixture, vortexed for 5 minutes in order to break the cells. Finally, 2 ml of water was added, tubes were vortexed for another 10 minutes and spun down for 5 minutes at 5000 rpm 4°C in order to separate the phases. Approximately 2 ml of aqueous phase was collected into a new tube, evaporated to dryness with (lyophilizer) and stored at −80°C. The culture samples taken before anaerobic incubation were treated the same way except washing step with NaCl.

A solution of d4-succinate (Sigma-Aldrich USA) was prepared to 53.1 mg/L in BG11 medium as an internal standard. Standards were prepared in the range 1–50 mg/L succinate in BG11 medium, these were filtered and diluted with the internal standard 4:1 by pipetting 5 μL internal standard solution into 20 μL succinate standards. Samples were diluted with internal standard in the same way. For determination of succinate levels in samples, supernatants and cell extracts were analysed with an Agilent 1100 HPLC connected to an Orbitrap LTQ Velos Pro (Thermo Fisher). The column a Hilicon iHILIC prototype (100 × 2.1 mm, 3.5 μm particle size), and mobile phase (70% 10 mM ammonium acetate, 30% acetonitrile) was pumped at 300 μL/min at 40 °C. Samples were injected at 1 μL. The detection was performed in SIM mode at 60,000 resolution, analyzing mass 117 and 121 sequentially. The resulting counts for the accurate masses of the two compounds were integrated and reported as ratio of succinate to the internal standard. The concentration of succinate in samples was determined by analysing the ratio of succinate: d4-succinate in the resulting solutions, and comparing with a linear calibration curve from the standards measured in triplicates.

### *SynLASPO* cloning and purification in *E. coli*

The *Synechocystis nadB* gene was amplified from genomic DNA as template by PCR according to standard protocols. The PCR-amplified sequence contained NdeI restriction site at 5’ and glycine-serine linker with Strep-tag placed at the C-terminus of the CDS, followed by a BamHI site at 3’, which were introduced with primer overhangs. The PCR fragment was cloned into the vector pET-11a (Novagen) using corresponding restriction sites. The resulting expression plasmid, pET-nadB, was used to transform *E. coli* BL21(DE3) cells. Recombinant protein expression was induced by adding 0.1mM of IPTG to the transformed *E. coli* cells grown to OD_600_ 0.4-0.6 and following overnight cultivation at 30°C. Cells were harvested by centrifugation at 7000rpm at 4°C for 10 minutes. Pellet was stored at −80°C.

Purification of *Syn*LASPO was carried out by Strep-Tactin affinity chromatography with StrepTrap™ XT (Cytiva) 5ml column on ÄKTA system (Cytiva) at 4 °C. For this, the pellet was thawed in PPB buffer (50 mM potassium phosphate buffer pH 8) containing 10% glycerol, 0.6 mg/ml lysozyme, 0.1 mg/ml DNAse, 0.1 mg/ml RNAse, 0.6 mg/ml inhibitor protease cocktail (G-Biosciences) and 20 μM FAD. After 30 minutes incubation on ice with lysozyme, the cell suspension was sonicated (Chemical Instruments AB, Model CV33) at 30% amplification, with 10 sec pulse and 20 sec rest intervals for 1 minute. The homogenate was then centrifuged at 50000 rpm for 40 min at 4°C. The obtained supernatant was loaded on the column pre-equilibrated with wash buffer (50 mM PPB pH 8, 10% glycerol). After column wash, the protein was eluted with 50mM biotin in wash buffer. The elution fraction was concentrated using Amicon Ultra-15 Centrifugal Filter Units MW 30 kDa, centrifuged at 5000 rpm for 20 min at 4 °C, frozen in liquid nitrogen and stored at −80°C. Protein concentration was determined with DC protein assay (Bio-Rad), using bovine serum albumin (Sigma) as a standard. The molecular weight of the purified protein was estimated by SDS-PAGE with Protein Plus prestained ladder (Thermo Fisher Scientific). UV–visible spectra of purified *Syn*Laspo were recorded using a Varian Cary 50 BIO spectrophotometer.

### Activity assays

Activity of *Syn*LASPO was determined by measuring the oxaloacetate derived from spontaneous hydrolysis of iminoaspartate [35]. The reaction mixture contained PPB with 10% glycerol, 100 μM FAD, various concentrations of aspartate and/or fumarate and 20 μM of purified protein. Anaerobic experiments were performed in a glovebox with 40 μM of purified protein. The mixture was incubated at 30°C, and 200 μL aliquots were taken at time points after the enzymatic reaction was initiated by addition of protein. Aliquots were quenched by addition of 100 μL 40% trichloroacetic acid followed by centrifugation at 12700 rpm for 10 minutes at 4°C degrees. Then, 250 μL of mixture was mixed with 100 μL of 5 mM 2,4-dinitrophenylhydrazine (DNPH) 5% HCl and incubated at 30°C for 30 minutes. Finally, 650 μL of 1M NaOH in 50% ethanol was added to the mixture which was left to stand for 10 minutes. Absorbance of the solution was read at 530nm, and the concentration of oxaloacetate formed during enzymatic reaction was determined based on an oxaloacetate calibration curve.

## Abbreviations

2-OG: 2-oxoglutarate
Cm: chloramphenicol
DNPH: 2,4-dinitrophenylhydrazine
*Escherichia coli*: *E. coli*
GABA: γ-aminobutyrate
Laspo: L-aspartate oxidase
SDH: succinate dehydrogenase
Sp: spectinomycin
*Synechocystis*: *Synechocystis* sp. PCC 6803
TCA cycle: tricarboxylic acids cycle

## Author contributions

KK and PL designed the study. KK and JK performed experiments. PL and CB supervised work in the project. KK, PL and CB analyzed the data. KK wrote the manuscript with PL, and CB and JH contributed to writing the manuscript. All authors read and approved the final manuscript.

## Acknowledgments

This work was supported by Formas (Grant No. 2016-01325), and by the Nord-Forsk NCoE program “NordAqua” (Project Number 82845).

## References

1. Cukalovic, A. & Stevens, C. V. (2008) Feasibility of production methods for succinic acid derivatives: a marriage of renewable resources and chemical technology, Biofuels, Bioproducts and Biorefining. 2, 505–529.

2. Yin, X., Li, J., Shin, H.-d., Du, G., Liu, L. & Chen, J. (2015) Metabolic engineering in the biotechnological production of organic acids in the tricarboxylic acid cycle of microorganisms: advances and prospects, Biotechnology advances. 33, 830–841.

3. Mock, M., Schmid, A. & Bühler, K. (2019) Photoautotrophic production of succinate via the oxidative branch of the tricarboxylic acid cycle influences glycogen accumulation in Synechocystis sp. PCC 6803, Algal Research. 43, 101645.

4. Mock, M., Schmid, A. & Bühler, K. (2020) Directed reaction engineering boosts succinate formation of Synechocystis sp. PCC 68O3_Δsll1625, Biotechnology Journal, 2000127.

5. Sengupta, A., Pritam, P., Jaiswal, D., Bandyopadhyay, A., Pakrasi, H. B. & Wangikar, P. P. (2020) Photosynthetic Co-Production of Succinate and Ethylene in A Fast-Growing Cyanobacterium, Synechococcus elongatus PCC 11801, Metabolites. 10.

6. Sengupta, S., Jaiswal, D., Sengupta, A., Shah, S., Gadagkar, S. & Wangikar, P. P. (2020) Metabolic engineering of a fast-growing cyanobacterium Synechococcus elongatus PCC 11801 for photoautotrophic production of succinic acid, Biotechnol Biofuels. 13, 89.

7. Lan, E. I. & Wei, C. T. (2016) Metabolic engineering of cyanobacteria for the photosynthetic production of succinate, Metab Eng. 38, 483–493.

8. Hasunuma, T., Matsuda, M. & Kondo, A. (2016) Improved sugar-free succinate production by Synechocystis sp. PCC 6803 following identification of the limiting steps in glycogen catabolism, Metab Eng Commun. 3, 130–141.

9. Hasunuma, T., Matsuda, M., Kato, Y., Vavricka, C. J. & Kondo, A. (2018) Temperature enhanced succinate production concurrent with increased central metabolism turnover in the cyanobacterium Synechocystis sp. PCC 6803, Metab Eng. 48, 109–120.

10. Osanai, T., Shirai, T., Iijima, H., Nakaya, Y., Okamoto, M., Kondo, A. & Hirai, M. Y. (2015) Genetic manipulation of a metabolic enzyme and a transcriptional regulator increasing succinate excretion from unicellular cyanobacterium, Front Microbiol. 6, 1064.

11. Ueda, S., Kawamura, Y., Iijima, H., Nakajima, M., Shirai, T., Okamoto, M., Kondo, A., Hirai, M. Y. & Osanai, T. (2016) Anionic metabolite biosynthesis enhanced by potassium under dark, anaerobic conditions in cyanobacteria, Sci Rep. 6, 32354.

12. Iijima, H., Watanabe, A., Sukigara, H., Iwazumi, K., Shirai, T., Kondo, A. & Osanai, T. (2021) Four-carbon dicarboxylic acid production through the reductive branch of the open cyanobacterial tricarboxylic acid cycle in Synechocystis sp. PCC 6803, Metabolic Engineering. 65, 88–98.

13. Xiong, W., Brune, D. & Vermaas, W. F. (2014) The γ-aminobutyric acid shunt contributes to closing the tricarboxylic acid cycle in S ynechocystis sp. PCC 6803, Molecular microbiology. 93, 786–796.

14. Zhang, S. & Bryant, D. A. (2011) The tricarboxylic acid cycle in cyanobacteria, Science. 334, 1551–1553.

15. Wan, N., DeLorenzo, D. M., He, L., You, L., Immethun, C. M., Wang, G., Baidoo, E. E., Hollinshead, W., Keasling, J. D. & Moon, T. S. (2017) Cyanobacterial carbon metabolism: Fluxome plasticity and oxygen dependence, Biotechnology and bioengineering. 114, 1593–1602.

16. Cooley, J. W. & Vermaas, W. F. (2001) Succinate dehydrogenase and other respiratory pathways in thylakoid membranes of Synechocystis sp. strain PCC 6803: capacity comparisons and physiological function, J Bacteriol. 183, 4251–8.

17. Osanai, T., Shirai, T., Iijima, H., Nakaya, Y., Okamoto, M., Kondo, A. & Hirai, M. Y. (2015) Genetic manipulation of a metabolic enzyme and a transcriptional regulator increasing succinate excretion from unicellular cyanobacterium, Frontiers in microbiology. 6, 1064.

18. Cooley, J. W., Howitt, C. A. & Vermaas, W. F. (2000) Succinate: quinol oxidoreductases in the cyanobacterium Synechocystis sp. strain PCC 6803: presence and function in metabolism and electron transport, Journal of Bacteriology. 182, 714–722.

19. Zhang, S., Qian, X., Chang, S., Dismukes, G. C. & Bryant, D. A. (2016) Natural and Synthetic Variants of the Tricarboxylic Acid Cycle in Cyanobacteria: Introduction of the GABA Shunt into Synechococcus sp. PCC 7002, Front Microbiol. 7, 1972.

20. Durall, C., Kukil, K., Hawkes, J. A., Albergati, A., Lindblad, P. & Lindberg, P. (2021) Production of succinate by engineered strains of Synechocystis PCC 6803 overexpressing phosphoenolpyruvate carboxylase and a glyoxylate shunt, Microbial cell factories. 20, 1–14.

21. Tedeschi, G., Negri, A., Mortarino, M., Ceciliani, F., Simonic, T., Faotto, L. & Ronchi, S. (1996) l-Aspartate Oxidase from Escherichia coli: II. Interaction with C4 Dicarboxylic Acids and Identification of a Novel l-Aspartate: Fumarate Oxidoreductase Activity, European journal of biochemistry. 239, 427–433.

22. Gerdes, S. Y., Kurnasov, O. V., Shatalin, K., Polanuyer, B., Sloutsky, R., Vonstein, V., Overbeek, R. & Osterman, A. L. (2006) Comparative genomics of NAD biosynthesis in cyanobacteria, J Bacteriol. 188, 3012–23.

23. Korshunov, S. & Imlay, J. A. (2010) Two sources of endogenous hydrogen peroxide in Escherichia coli, Mol Microbiol. 75, 1389–401.

24. Englund, E., Andersen-Ranberg, J., Miao, R., Hamberger, B. r. & Lindberg, P. (2015) Metabolic engineering of Synechocystis sp. PCC 6803 for production of the plant diterpenoid manoyl oxide, ACS synthetic biology. 4, 1270–1278.

25. Liu, X., Miao, R., Lindberg, P. & Lindblad, P. (2019) Modular engineering for efficient photosynthetic biosynthesis of 1-butanol from CO 2 in cyanobacteria, Energy & Environmental Science. 12, 2765–2777.

26. Bossi, R. T., Negri, A., Tedeschi, G. & Mattevi, A. (2002) Structure of FAD-bound L-aspartate oxidase: insight into substrate specificity and catalysis, Biochemistry. 41, 3018–3024.

27. Mortarino, M., Negri, A., Tedeschi, G., Simonic, T., Duga, S., Gassen, H. G. & Ronchi, S. (1996) L-Aspartate Oxidase from Escherichia coli: I. Characterization of Coenzyme Binding and Product Inhibition, European journal of biochemistry. 239, 418–426.

28. Bifulco, D., Pollegioni, L., Tessaro, D., Servi, S. & Molla, G. (2013) A thermostable L-aspartate oxidase: a new tool for biotechnological applications, Applied microbiology and biotechnology. 97, 7285–7295.

29. Chapman, S. K. & Reid, G. A. (1999) Flavoprotein protocols, Springer Science & Business Media.

30. Yamada, R.-h., Nagasaki, H., Wakabayashi, Y. & Iwashima, A. (1988) Presence of D-aspartate oxidase in rat liver and mouse tissues, Biochimica et Biophysica Acta (BBA)-General Subjects. 965, 202–205.

31. Mattevi, A., Tedeschi, G., Bacchella, L., Coda, A., Negri, A. & Ronchi, S. (1999) Structure ofL-aspartate oxidase: implications for the succinate dehydrogenase/fumarate reductase oxidoreductase family, Structure. 7, 745–756.

32. Hao, J., Petriacq, P., de Bont, L., Hodges, M. & Gakiere, B. (2018) Characterization of l-aspartate oxidase from Arabidopsis thaliana, Plant Sci. 271, 133–142.

33. Hurley, J. K., Morales, R., Martinez-Júlvez, M., Brodie, T. B., Medina, M., Gómez-Moreno, C. & Tollin, G. (2002) Structure–function relationships in Anabaena ferredoxin/ferredoxin: NADP+ reductase electron transfer: insights from site-directed mutagenesis, transient absorption spectroscopy and X-ray crystallography, Biochimica et Biophysica Acta (BBA)-Bioenergetics. 1554, 5–21.

34. Santos, A. R. S., Gerhardt, E. C. M., Parize, E., Pedrosa, F. O., Steffens, M. B. R., Chubatsu, L. S., Souza, E. M., Passaglia, L. M. P., Sant’Anna, F. H. & de Souza, G. A. (2020) NAD+ biosynthesis in bacteria is controlled by global carbon/nitrogen levels via PII signaling, Journal of Biological Chemistry. 295, 6165–6176.

35. Marinoni, I., Nonnis, S., Monteferrante, C., Heathcote, P., Härtig, E., Böttger, L. H., Trautwein, A. X., Negri, A., Albertini, A. M. & Tedeschi, G. (2008) Characterization of L-aspartate oxidase and quinolinate synthase from Bacillus subtilis, The FEBS journal. 275, 5090–5107.

36. Ollagnier-de Choudens, S., Loiseau, L., Sanakis, Y., Barras, F. & Fontecave, M. (2005) Quinolinate synthetase, an iron–sulfur enzyme in NAD biosynthesis, FEBS letters. 579, 3737–3743.

37. Saunders, A. H., Griffiths, A. E., Lee, K.-H., Cicchillo, R. M., Tu, L., Stromberg, J. A., Krebs, C. & Booker, S. J. (2008) Characterization of quinolinate synthases from Escherichia coli, Mycobacterium tuberculosis, and Pyrococcus horikoshii indicates that [4Fe-4S] clusters are common cofactors throughout this class of enzymes, Biochemistry. 47, 10999–11012.

38. Stanier, R. & Cohen-Bazire, G. (1977) Phototrophic prokaryotes: the cyanobacteria, Annual review of microbiology. 31, 225–274.

39. Heidorn, T., Camsund, D., Huang, H.-H., Lindberg, P., Oliveira, P., Stensjö, K. & Lindblad, P. (2011) Synthetic biology in cyanobacteria: engineering and analyzing novel functions in Methods in enzymology pp. 539–579, Elsevier.

40. Prasannan, C. B., Jaiswal, D., Davis, R. & Wangikar, P. P. (2018) An improved method for extraction of polar and charged metabolites from cyanobacteria, PLoS One. 13, e0204273.

