## Supplementary information for "L-Aspartate oxidase provides new insights into fumarate reduction in anaerobic darkness in *Synechocystis* sp. PCC6803"

Supporting information

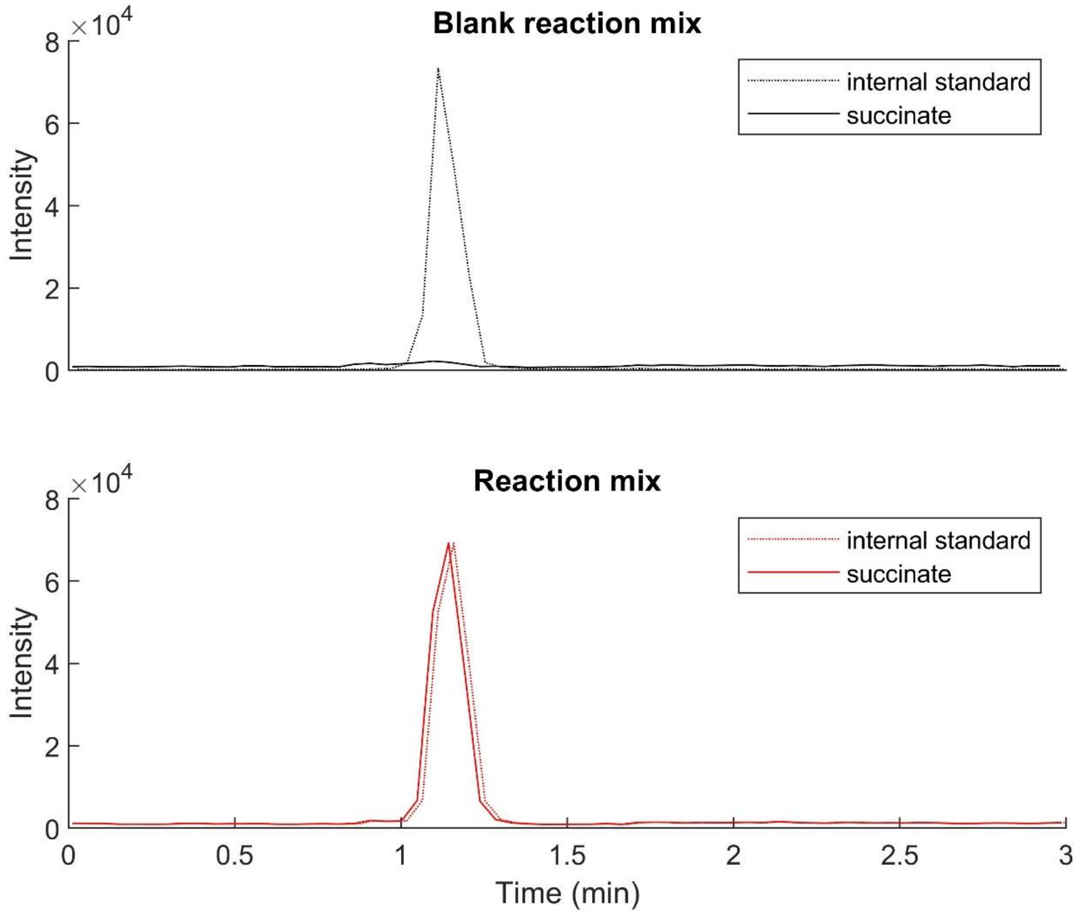


Figure S1. Succinate peaks observed during anaerobic experiments using isocratic hydrophilic interaction chromatography coupled to mass spectrometry (for method see method section of main paper). Peak area is proportional to concentration, and both analytical mixtures contained 10 ppm of a deuterated internal standard (d4-succinate). *Syn*Laspo enzyme was absent in blank reaction mix; both reaction mixes contained 1mM fumarate.
